# UVA hyperspectral light-sheet microscopy for volumetric metabolic imaging: application to pre-implantation embryo development

**DOI:** 10.1101/2023.06.21.545939

**Authors:** J. Morizet, D. Chow, P. Wijesinghe, E. Schartner, G. Dwapanyin, N. Dubost, G. D. Bruce, E. Anckaert, K. Dunning, K. Dholakia

## Abstract

Cellular metabolism is a key regulator of energetics, cell growth, regeneration and homeostasis. Spatially mapping the heterogeneity of cellular metabolic activity is of great importance for unraveling the overall cell and tissue health. In this regard, imaging the endogenous metabolic co-factors NAD(P)H and FAD with sub-cellular resolution and in a non-invasive manner would be useful to determine tissue and cell viability in a clinical environment, but practical use is limited by current imaging techniques. In this article, we demonstrate the use of phasor-based hyperspectral light-sheet (HS-LS) microscopy using a single UVA excitation wavelength as a route to mapping metabolism in three dimensions. We show that excitation solely at a UVA wavelength of 375 nm can simultaneously excite NAD(P)H and FAD autofluorescence, while their relative contributions can be readily quantified using a hardware-based spectral phasor analysis. We demonstrate the potential of our HS-LS system by capturing dynamic changes in metabolic activity during pre-implantation embryo development. To validate our approach, we delineate metabolic changes during pre-implantation embryo development from volumetric maps of metabolic activity. Importantly, our approach overcomes the need for multiple excitation wavelengths, two-photon imaging or significant post-processing of data, paving the way towards clinical translation, such as in situ, non-invasive assessment of embryo viability.

## Introduction

Metabolism is central to fulfilling biological functions of living cells, underpinning key processes in development, regeneration and homeostasis.^1^ The quantification of metabolic activity has, thus, been an important endeavour to gain insight into live cell and tissue health. This has been accelerated by recent findings that have highlighted its potential as a diagnostic marker for cancer^2–5^ and neurodegenerative disease.^6,7^ Further, metabolic activity has been shown to be a reliable indicator of treatment efficacy in cancer organoids^8,9^ and an indicator of viability in tissue engineering^10,11^ and for the developing embryo.^12–14^ Advances in photonics have been at the forefront of research in cellular metabolism because they offer a route to monitor metabolic activity in situ via fluorescence. To date, many volumetric imaging techniques with sub-cellular resolution have been restricted in use solely in a research capacity. This is because they fail to meet the imaging speed and permissible phototoxicity required for many clinical applications. Furthermore, they are often inaccessible outside a research environment due to the complexity, availability and cost of the required instrumentation equipment.^15^ Enabling adoption for routine biological and clinical assessment requires the development of non-invasive, high resolution label-free imaging techniques with minimal technical complexity. A powerful example of this demand can be found in clinical in vitro fertilization (IVF) procedures, where metabolic activity can reveal embryos with higher developmental potential.^16^ Non-invasive optical imaging techniques have the potential to improve embryo selection and revitalize in the presently stagnant 30% success rate of a live birth.^17,18^

Cellular metabolic activity can be quantified non-invasively by monitoring autofluorescence from endogenous metabolic co-enzymes nicotinamide adenine dinucleotide (phosphate) (NAD(P)H) and flavin adenine dinucleotide (FAD). A common metric to quantify metabolism is the redox ratio (RR), which may be defined as the normalized ratio of the molecular concentrations of the two metabolites.^19^ Ratiometric quantification of the normalized ratio of both metabolites has been assessed spatially using several imaging modalities, such as epifluorescence microscopy,^14^ confocal microscopy,^20,21^ multiphoton microscopy^22^ and fluorescence lifetime imaging microscopy (FLIM).^2^ The major challenge of all these modalities is the accurate unmixing and quantification of the fluorescent species. This is typically addressed by employing multiple excitation wavelengths and spectral detection using appropriate emission bands commensurate with the fluorescence profiles of NAD(P)H and FAD. Such measurements have enabled the assessment of metabolic activity in numerous pathological conditions including cancer^2–5^ and neurological diseases.^6,7^

Some recent efforts have been made to quantify metabolic activity using light-sheet microscopy (LSM). LSM is a powerful volumetric, rapid imaging modality with sub-micron resolution that is well-known for its minimal photodamage due to its geometry and fast image acquisition.^23^ In this regard, LSM contrasts other imaging modes, such as FLIM, which have long integration times^24^ that impedes their use for high-throughput viability screening and long-term volumetric metabolic imaging. There have been few studies to date based on light-sheet for metabolic imaging. Favreau et al.^25^ sequentially excite NAD(P)H and FAD metabolic co-enzymes using two wavelengths of 405 nm and 488 nm, respectively, to assess the response of colorectal cancer organoids to treatments. Dual-wavelength excitation requires precise co-alignment of the illumination beams and the inherent longitudinal chromatic aberration and dispersion may lead to disparate fields of view and focal positions for each wavelength. Spatial artifacts in RR assessment may also arise from differences in scattering and attenuation properties between the two wavelengths as a function of depth. Furthermore, the subsequent impact of the different laser power at the two wavelengths and detector sensitivity for two collection channels on the fluorescence intensity levels collected for NAD(P)H and FAD typically requires correction in a post-processing step via rigorous calibration using reference solutions. Recently, Hedde et al.^26^ demonstrated that two-photon LSM at 740 nm coupled with a new hyperspectral detection scaheme can distinguish a variation in the metabolic redox ratio between the crypt and lumen of mouse colon. While promising, multiphoton LSM requires a costly and high-pulse-energy laser source, thus, is rarely compatible with a clinical setting.

In contrast to these studies, we present a light-sheet microscopy system that employs a single-wavelength, one-photon excitation in the UVA range at 375 nm. Single-wavelength excitation is enabled by the sensitive hardware-based unmixing for accurate, fast metabolic mapping in all three dimensions and without additional calibration steps. Our study reveals both numerically and experimentally that, in the case of NAD(P)H and FAD co-excitation, performing standard bandpass filtering of the emission spectra fails to accurately quantify RR, which has been a major limitation of single-wavelength excitation. This is further complicated by the lack of standardized bandpass filters, which prevents direct comparison between studies. To circumvent this issue, here we adopt a hardware-based spectral phasor analysis of fluorescence ^26^ to quantify RR by sole use of two cosine and sine transmission filters in our LSM approach.

We demonstrate the utility of our approach by performing metabolic imaging of pre-implantation embryos, which is a burgeoning need in clinical embryology.^13^ We clearly delineate changes in metabolic state within key stages of embryo development from the 2-cell up to the blastocyst-stage. Mapping metabolic activity in a volume with minimal photo-damage is necessary to capture embryo-level metabolic heterogeneity, which is known to be crucial in the case of embryos with cells containing an unexpected number of chromosomes (mosaicism).^14^ Collectively, we propose that our rapid 3D imaging technique offers minimal technical complexity and can capture metabolic activity, opening up new opportunities for rapid, high-throughput metabolic mapping of 3D samples in clinical contexts.

## Results and discussion

The key challenge of single-wavelength excitation for RR monitoring is the ability to co-excite NAD(P)H and FAD fluorophores, while unmixing their relative contributions to the total emitted intensity with high sensitivity. Several definitions of RR are generally proposed in the literature, and we adopt here the convention where RR is defined as the ratio of FAD intensity divided by the sum of NAD(P)H and FAD intensities, as follows: *I_F AD_/*(*I_NAD_*_(_*_P_* _)_*_H_* + *I_F AD_*).^19^ Importantly, this parameterisation of RR demands that the recorded intensities reflect the underlying concentration of the metabolic co-enzymes. Thus, it is imperative that the product of the molar absorbance, quantum yield and integrated probability density of each metabolite, as well as the detection efficiency of the imaging method is relatively equivalent for NAD(P)H and FAD, as detailed in the *Section S1 of supplementary information*. While most of these factors are intrinsic properties of the fluorophores or the setup and cannot be tuned, the absorbance depends on the wavelength used for the excitation, which therefore needs to be carefully selected. Figure 1a shows that one-photon NAD(P)H fluorescence absorption and emission maxima are approximately at 350 nm and 460 nm, respectively, while one-photon FAD fluorescence absorption maxima can be found at 350 nm and 450 nm, with a fluorescent emission maxima at 535 nm, as reported in the literature.^27,28^ In this regard, the illumination wavelength is a key parameter that can be adjusted to achieve a desired equivalence in the NAD(P)H and FAD absorbance. An absorbance of the same magnitude for NAD(P)H and FAD can be obtained with an illumination in the UVA range (315–400 nm), which ensures an approximate linearity between RR and co-enzyme concentration (derived in detail in the *Section S1 of supplementary information*).

**Figure 1:**
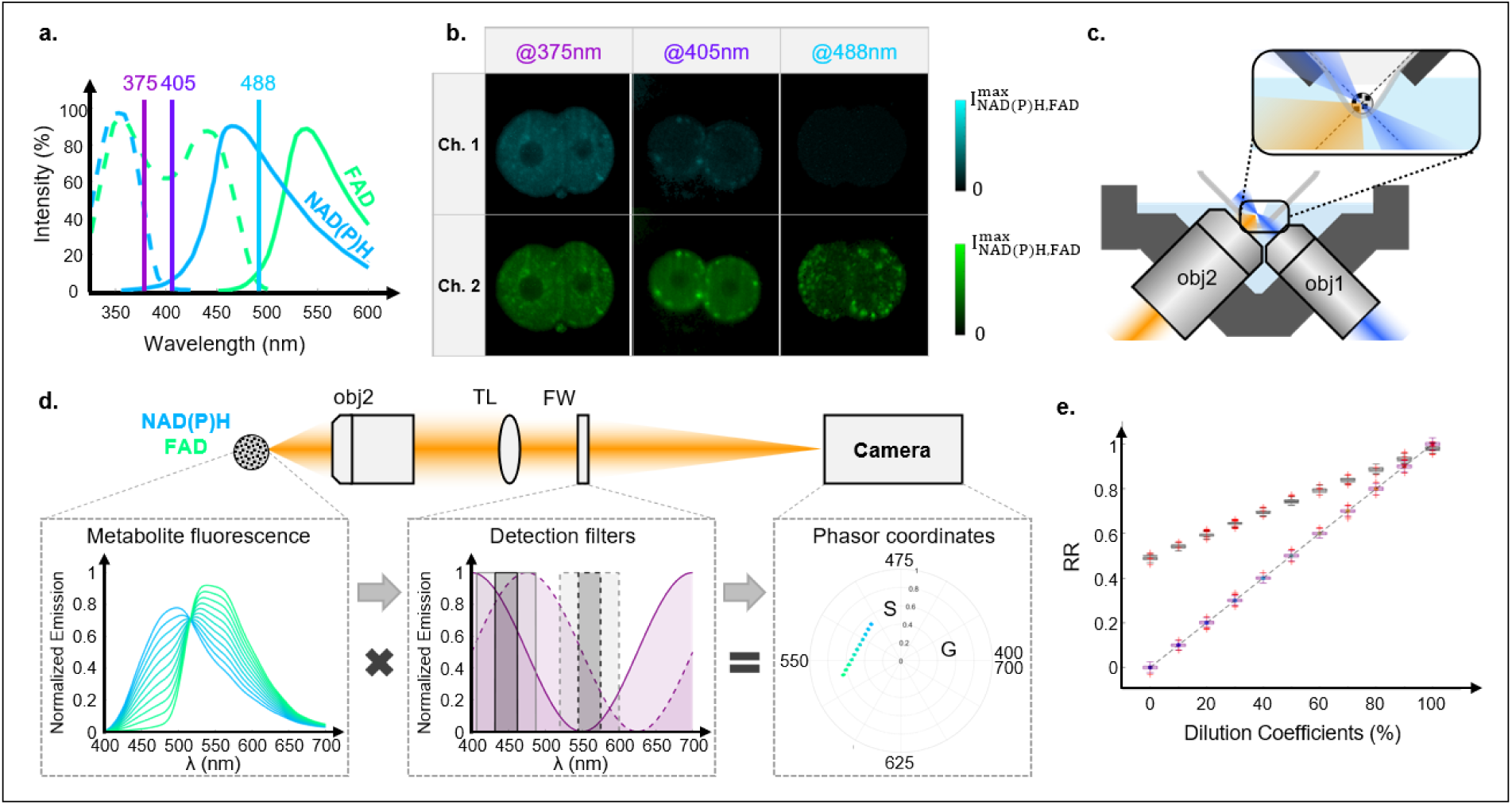
(a) Absorption (dashed line) and emission (solid line) spectra of NAD(P)H (blue) and FAD (green) metabolites reported in^27,28^ with additional markers for *λ_exc_* values of 375, 405, 488 nm. (b) Representative intensity images collected in the channel Ch1 (447/60nm) and Ch2 (560/94nm) using 375, 405, 488 nm excitations. (c) Open-top light-sheet configuration. The sample is placed on a custom-designed V-mount. This is illuminated from the right through the illuminating objective (obj1) and fluorescent light is collected by the detecting objective (obj2) before passing through the detection path shown in (a). (d) Schematic of the hyperspectral detection where fluorescence signals are converted to non-normalized phasor coordinates (G,S) using cosine/sine transmission filters. (e) Comparison between RR evaluated using spectral phasor-based detection (purple) and bandpass filters (grey).

To understand how the choice of the excitation wavelength translates in terms of intensity contrast, we imaged mouse embryos and compared the contrast obtained in two spectral channels with different bandpass filters (Ch1: 447/60 nm; Ch2: 560/94 nm) and with illuminations at 375, 405 and 488 nm (see Figure 1b). Unsurprisingly, excitation at 488 nm provides contrast only in Ch2 (commensurate with FAD emission) with clear visualization of bright domains within the cells, likely due to the presence of enriched FAD within mitochondria.^29^ In comparison, excitation at 405 nm provides a low, yet relatively uniform contrast in Ch1 and a high and moderately uniform contrast in Ch2. This can be understood as a co-excitation of NAD(P)H and FAD, with the total intensity dominated by FAD fluorescence along with a small contribution from NAD(P)H leaking in Ch2. Lastly, excitation at 375 nm provides higher contrast in Ch1 compared to previous excitations, indicating higher NAD(P)H intensity, while FAD is co-excited. Therefore, to co-excite FAD and NAD(P)H simultaneously, we use a continuous laser source at 375 nm, which is an easily accessible source. Notably, UV excitation has been used in early spectroscopy studies to isolate NAD(P)H contribution,^20^ as well as in recent studies for NAD(P)H and FAD co-excitation using an illumination wavelength of 375 nm.^30,31^ However, it was rarely used for imaging due to the lack of sectioning capability of common one-photon fluorescence imaging techniques, as confocal microscopy, combined with potential photodamage caused by a high light dose in addition to long exposure times.

Based on this wavelength selection, we experimentally assessed metabolic activity in mouse embryos using a custom-built open-top light-sheet geometry with Gaussian-beam illumination as illustrated in Figure 1c and detailed in *Section S2 of supplementary information*. We incorporated two spectral detection schemes, including: (i) a set of bandpass filters, and (ii) filters with specific sine/cosine transmission following a phasor-based hyperspectral detection method proposed by Hedde et al.^26^ (see Figure 1d). The spectral phasor analysis was originally introduced by Fereidouni et al.^32^ to provide a fit-free analysis of spectral data. This consists of a post-processing step that transforms the fluorescence spectra into their *n^th^* order Fourier spectral components: 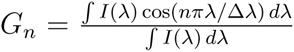 and 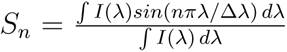, where Δ*λ* = *λ_max_ − λ_min_*. *G_n_* and *S_n_* can be represented as orthogonal components on a phasor plot. For simplicity, spectral analysis can be limited to the 1*^st^* Fourier order (*n* = 1), which describes the spectral barycenter of the emission. A direct hardware transformation of the fluorescence into (G,S) phasor coordinates has been proposed recently by Hedde et al.^26^ This hyperspectral approach is based on the addition of two spectral filters into the detection path with *T*_cos_(*λ*) and *T*_sin_(*λ*) transmission profiles that follow a single cosine and sine period respectively in the 400–700 nm wavelength range. Each filter converts the sample-emitted fluorescent light directly into the Fourier components, G and S, which are sequentially projected onto the camera by means of a tube lens in our setup (see Figure 1d).

The quantification of RR values is a typical challenge in the case of co-enzyme co-excitation. The common approach to estimate RR values is to use two bandpass filters with bandwidths centered on the maxima of NAD(P)H and FAD emission, i.e.: 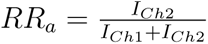, where Ch1 collects photons in the channel centered at 450 nm and Ch2 detects photons in the channel centered at 550 nm sequentially from two separate excitation wavelengths. Figure 1e shows *RR_a_* values derived numerically from normalized emission spectra of mixed solutions of NAD(P)H and FAD with a linear dilution coefficient. When using single-wavelength excitation, *RR_a_* values were found to increase from around 0.5 to 1 for filters with bandwidths of 30 and 80 nm when the dilution coefficient varies from 0% (NAD(P)H only) to 100% (FAD only). This large discrepancy between *RR_a_* values compared to the expected values— which should vary between 0 and 1 for a similar distribution of the dilution coefficient—is due to the cross-talk between NAD(P)H and FAD in the collected channels. As a result, a substantial contribution of NAD(P)H fluorescence in the FAD detection channel translates to an incorrect parameterisation: 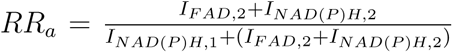, where the subscript denotes the channel in which the fluorescence is collected.

Next, we compared numerical RR values quantified with the conventional method with the values resulting from spectral phasor analysis. Figure 1d shows spectral phasor coordinates of the solutions under study. An interesting property of this approach, known as the linear addition law, dictates that mixed contributions lie linearly along a line that joins the coordinates of the pure fluorescent species. As derived in detail in *Section S2 in supplementary material*, RR can be calculated as 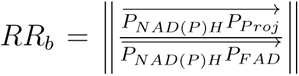, i.e., the distance between the NAD(P)H coordinate and any projected coordinate normalized by the distance between the NAD(P)H-FAD coordinates. Figure 1e illustrates that, in contrast to RR values assessed for bandpass filters, *RR_b_* values do not suffer from cross talk and accurately represent the mixing ratio from 0 and 1. As noted above, the RR evaluated with the hyperspectral approach provides a value that accurately estimates 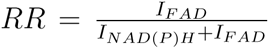 and offers a clear advantage by providing an accurate RR value consistent with the original definition. In addition, the precision of both methods was also investigated in *Section S1 of supplementary material* by calculating the ratio of the normalized RR standard deviation to the RR gradient. The precision of the spectral phasor analysis was found to be superior to conventional bandpass detection. It is also worth noting that for wide spectral bandpass filters, the precision approaches that of spectral phasor analyses due to the larger number of photons collected. In *Section S1 of supplementary material*, we have further explored the effect of non-normalization for pure solutions intensities, which can occur when the two absorbance associated with the illumination wavelength are not of the same order of magnitude. As a result, we observe a loss of linearity in RR values, however, the hyperspectral method remains superior to bandpass filters.

Figure 2a–g illustrates the recovery of metabolic information on a single cross-sectional plane of a blastocyst-stage embryo. The imaged intensities, corresponding to phasor coordinates (G,S), were background corrected and normalized for each pixel (see the Methods section for details). Since any change in embryo metabolism is likely to result in a translation in the phasor coordinates along the NAD(P)H-FAD trajectory, the phasor coordinates of pure NAD(P)H and FAD solutions were experimentally characterized (Figure 2a), then phasor coordinates in each pixel of the image were orthogonally projected along the NAD(P)H-FAD trajectory (Figure 2b). For visualization, *RR_b_* has been color-coded using the color bar shown in Figure 2c. Figures 2d–f show the total autofluorescence intensity, the phasor plot of all pixels and the distribution of *RR_b_* in the embryo, respectively. The distribution of the *RR_b_* data along the NAD(P)H-FAD trajectory was further summarized as a histogram (Figures 2c,g).

**Figure 2:**
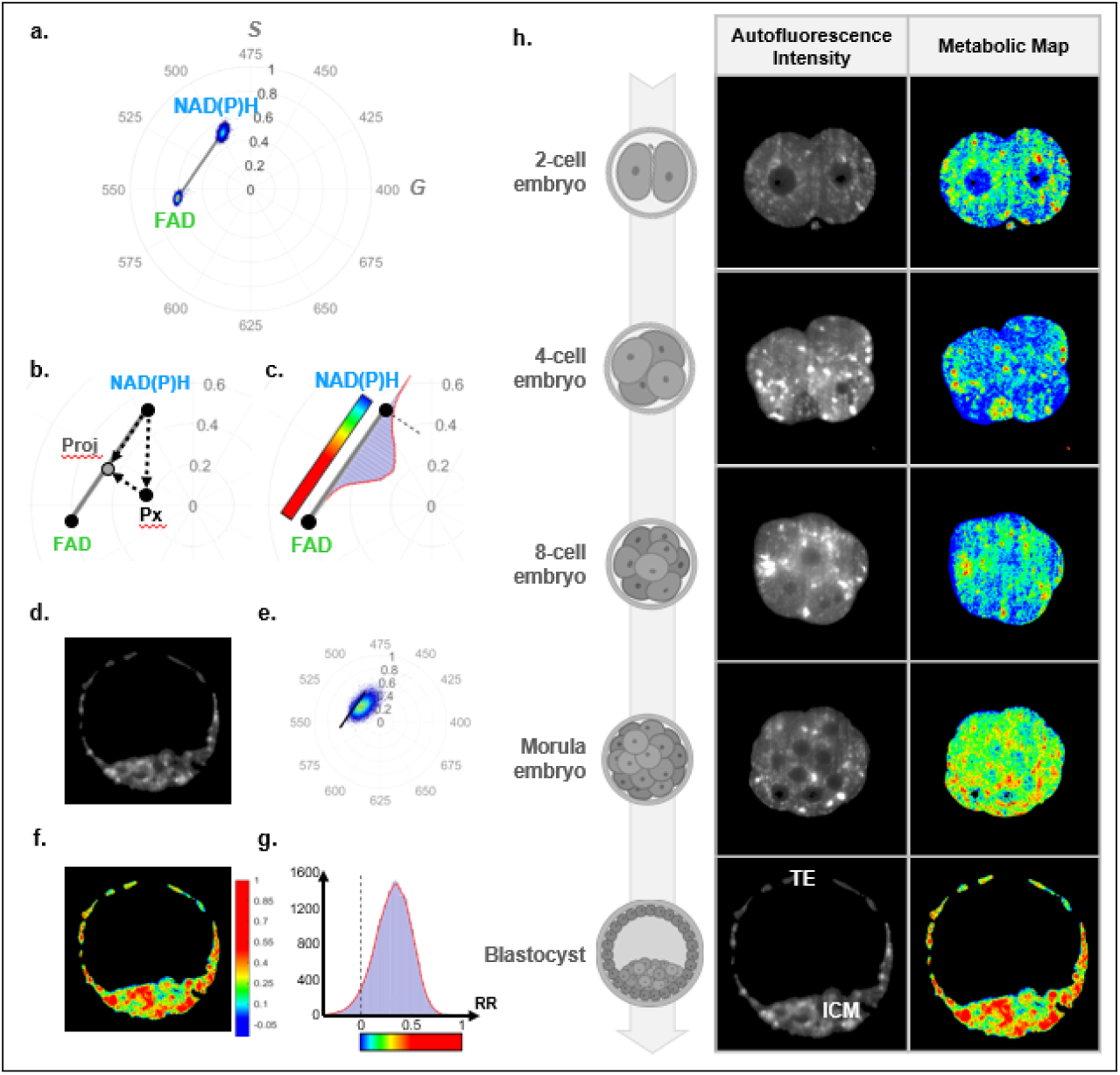
Phasor plots with (a) experimentally measured coordinates of pure NAD(P)H and FAD solutions, (b) RR assessment method based on orthogonal projection of the normalized phasor coordinates (G, S) along the NAD(P)H-FAD trajectory and (c) the color map encoding RR values for the metabolic maps. Example of (d) autofluorescence intensity map, (e) phasor coordinates represented on a polar plot, (f) metabolic map and (g) histogram plot generated by the data analysis pipeline applied on a single imaged plane acquired on a blastocyst-stage embryo. (h) Schematic representation of 2-, 4-, 8-cell, morula, and blastocyst-stage embryo with their corresponding autofluorescence intensity images and metabolic map. ICM and TE depict the subpopulation of cells - inner cell mass and trophectoderm respectively - found within a blastocyst-stage embryo.

Using our approach, we analysed 2D and 3D hyperspectral acquisitions of embryos throughout stages of its development. Figure 2h presents the autofluorescence intensity and the corresponding metabolic maps of a single cross-section acquired at different embryo developmental stages from the 2-cell up to the blastocyst-stage. The intensity contrast reveals a morphological evolution of an embryo over time. The 2-cell embryo undergoes cellular division (mitosis) which ultimately transforms it into a densely packed cluster of cells (successively 4-cell, 8-cell and morula). Subsequently, this population of cells undergoes cellular differentiation where pluripotent embryonic cells within an embryo commit to forming either the inner cell mass (ICM, fetal-lineage) or the trophectoderm (TE, placental-lineage) following the formation of a fluid-filled cavity known as the blastocoel cavity. The colorimetric changes in the metabolic maps suggest the possibility of using metabolic variations over time for monitoring embryo development, which we further study and discuss below.

Figure 3 shows examples of 3D visualizations of the morphological and metabolic content of embryos with three side views at the 4-cell (Figure 3a), morula (Figure 3b) and blastocyst-stage (Figure 3c). In addition, we present a 3D movie which represents a metabolic map of these embryos in the *supplementary material*. Interestingly, the metabolic map shows heterogeneity throughout the embryos, with higher RRs confined to small domains which are likely to be cell mitochondria. This highlights the potential for 3D reconstruction to be used in further studies to conjointly investigate morphological and metabolic properties.

**Figure 3:**
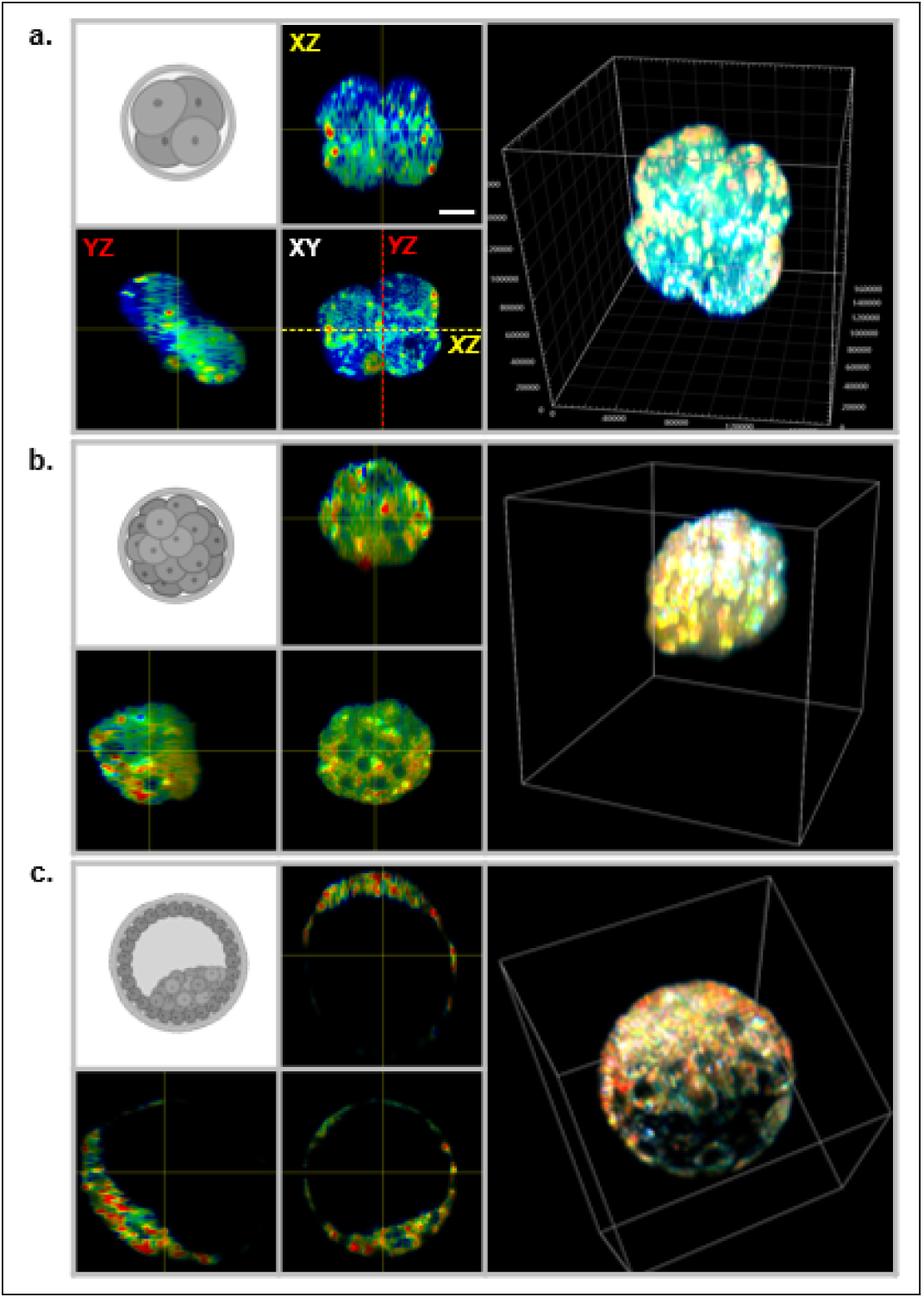
3D reconstruction of embryo metabolic maps. Gray schematic diagrams depict the aspect of embryos in (a) a 4-cell embryo, (b) a morula and (c) a blastocyst stage embryo. XY, XZ and YZ views along with 3D reconstruction of the metabolic maps are shown for these different stages of development. Scale bar = 20*µ*m

To assess safety for embryos throughout imaging, illuminated embryos were placed in the incubator immediately after imaging and their mortality rates were compared to a non-illuminated (control) group. We found no statistical difference in the number of embryos that developed to the blastocyst-stage between the two groups (see *Section S3 in supplementary material*). The illumination levels used here, therefore, likely have no measured effect on embryonic development from temporal development and morphological considerations when using rapid light-sheet imaging with an excitation at 375 nm.

Finally, we investigated the capability to quantify changes in metabolic activity during development. This was achieved with a 3D analysis on imaging data acquired on embryos throughout development. The mean RR was determined by averaging RR over all pixels of XYZ image stack generated for each embryo. Further, we have acquired RR values using the conventional method using bandpass filters during the same experiment for comparison, i.e., *RR_a_* = *I_Ch_*_2_*/*(*I_Ch_*_1_ + *I_Ch_*_2_). Figures 4a and b show the temporal evolution of the two metabolic variables obtained with both methods.

**Figure 4:**
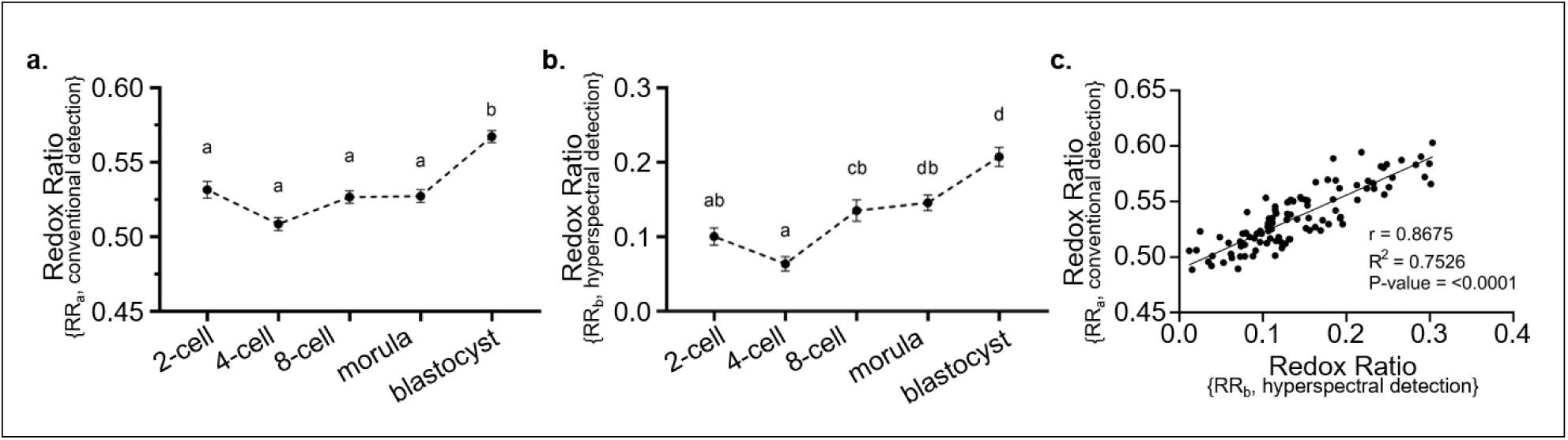
RR computed following volumetric imaging of embryos through development. Embryos were excited with a 375 nm laser and signals collected via (a) conventional method with 2 bandpass filters and (b) hyperspectral detection. Data are presented as mean *±* SEM, n=11-26 embryos per developmental stage, from 3 independent experimental replicates. Data were analyzed using the Kruskal-Wallis test with Dunn’s multiple comparisons test. Different superscripts indicate statistical significance (*P <*0.05) between developmental stages. (c) Correlation graph between RR values assessed using conventional and hyperspectral detection.

It is clear that the range of RR values obtained with bandpass filtering differs from the one using the phasor-based approach, with *RR_a_* found to be between 0.51 and 0.57 and *RR_b_* values found to be between 0.06 and 0.21 respectively. This discrepancy is predicted and discussed in Figure 1e, and manifests due to the cross-talk of co-excited NAD(P)H and FAD emission spectra in the detection channels. This results in a lack of accuracy in *RR_a_* assessment with the bandpass filter approach, leading to an overestimation of the RR.

In addition, the conventional approach not show statistically significant variations during early embryo development (Figure 4a; P*>*0.05), with a statistically significant higher *RR_a_* value found only at the blastocyst-stage embryos. In comparison, the hyperspectral method showed a gradual and statistically significant increase in *RR_b_* values from the 8-cell up to the blastocyst-stage (Figure 4b; P*>*0.05) but with no significant changes in *RR_b_* at earlier stages of development. The low and statistically unchanged values reported for both metabolic metrics between 2- and 8-cell stages indicate that embryo metabolism remains low in the early stages of pre-implantation embryonic development.

The following increase in RR consistently observed with these two methods reflects a change in embryo metabolism characterized by a higher metabolic activity. This result corroborates the shift from oxidative phosphorylation to glycolysis as reported in the literature,^33^ which coincides with an increased bio-energetic demand for cellular proliferation and differentiation as well as changes in the bioavailability of energy substrate in its microenvironment.^33^ Further, we observe a greater precision in assessing metabolic changes using the hyperspectral method throughout development. We see shifts from low to high metabolism as embryos develop, which coincide with the timing of embryonic genome activation, which occurs at the 2- to 4-cell stage in mice^34^ and the shift in metabolic pathway utilised as mentioned above.

While metabolic studies using volumetric imaging with subcellular resolution have been limited to a research context because of low throughput in terms of imaging speed and permissible photodamage, as well as have demonstrated being difficult access outside a research environment due to the equipment required, we showed here the benefits of UVA single excitation light-sheet microscopy combined with the phasor-based approach for mouse embryos imaging. The use of HS-LS harnesses the intrinsic advantages of light-sheet microscopy with rapid 3D imaging and minimal sample illumination. Further, single wavelength one-photon excitation at 375 nm allows NAD(P)H and FAD co-excitation without the need for costly and difficult-to-access sources, enabling further dissemination of this technology. The simplicity of installation is a major asset from an instrumental perspective for a device designed for a clinical environment, unlike multi-wavelength instruments. This helps to overcome issues encountered with a dual excitation such as co-alignment of illumination beams, dissimilar fields of view and focus positions, and spectral properties of the scattering inducing differences in attenuation as a function of the illumination wavelengths. The facile implementation of a single wavelength for excitation was made possible by spectral phasor-based filtering, which enabled RR assessment with superior accuracy in the case of NAD(P)H and FAD co-excitation compared to bandpass filter methods. Further, adapting miniaturized light-sheet microscopy^35,36^ with an illumination in the UVA range at 375 nm combined with a phasor-based hyperspectral detection would enable the system to be portable for studies in a clinical environment. Although our initial demonstration is limited to assessing metabolic changes in embryos at different stages of development, this study paves the way for metabolic assessment in mouse embryos to evaluate their viability for implantation in a clinical context. Future studies could consider the transfer of embryos into pseudo-pregnant female mice following metabolic assessment using HS-LS to assess for potential pregnancy and birth complications associated with exposure to UVA wavelength during pre-implantation embryo development. Also, combined with microfluidic devices, this approach also promises high-throughput metabolic assessment in a large variety of models ranging from embryos to organoids to evaluate viability or treatment efficiency.

## Conclusions

We have demonstrated an original single 375 nm wavelength phasor-based hyperspectral light-sheet microscope for volumetric metabolic mapping. The autofluorescence signal for NAD(P)H and FAD were collected using both conventional detection with two separate band-pass filters and hardware-based hyperspectral detection following excitation at a wavelength of 375 nm. Hyperspectral detection was shown to be more precise and accurate to changes in embryo metabolism through development compared to conventional detection of NAD(P)H and FAD signals. The method promises accurate 3D mapping of cellular metabolism, which can benefit areas such as analyses of organoids, cell spheroids and pre-implantation embryo. The facile implementation of this system opens up opportunities for high-throughput, accessible imaging of embryos to study the temporal evolution of metabolism in clinical environments.

## Methods

### Set-up

Imaging was performed using a custom-built open-top virtual light-sheet microscope with three continuous-wave laser sources emitting in the UVA-visible range (see *Section S1 of Supplement1*). The first UVA beam is delivered in free space by a 375 nm laser (*Stradus 375-60, Vortran*). It is magnified with a first telescope (*f*_1_ = 40*mm*, *f*_2_ = 75*mm*) and spatially filtered using a pinhole (*ϕ* = 30*µm*) inserted between the two lenses. The second and third beams originate from a pair of 405 nm and 488 nm fiber-coupled lasers (*Obis FP405LX, FP488LX, OBIS LX/LS Laser Box*) and are coupled into a single fiber via a combiner (*OBIS Galaxy Beam Combiner System, Coherent*), then collimated in free space by a *f*_3_ = 25*mm* lens. To precompensate for any longitudinal chromatic aberration accumulated along the illumination path to the imaging plane, the 405 nm and 488 nm beams were separated again using a dichroic (D2, *DMLP425R, Thorlabs*) beam splitter and relayed through 1:1 telescopes (*f*_4_ = 50*mm*) placed in their respective optical paths, thereby finely correcting for divergence and obtaining similar focal planes. A similar dichroic (D3) beam splitter then recombined the 405 nm beam with the 488 nm beam, before a third dichroic beam splitter (D1, *FF389-Di01-25X36X1.5, Semrock*) was used to recombine them with the 375 nm laser beam. A 1D rotation galvo-mirror (*GVS201, Thorlabs*) was placed conjugate with the back pupil plane of the water-dipping 10X illumination objective (obj1, *UMPLFLN10XW, Olympus*) via an expansion telescope (*f*_5_ = 50*mm*, *f*_6_ = 75*mm*) to laterally scan the Gaussian beam in the field of view. Both the illumination objective and the 40X collection objective (obj2, *CFI Apo NIR 40X W, Nikon*) were inserted at 45*^◦^* from the horizontal plane into a custom-made mount to achieve the open-top geometry. The embryos were placed in a custom-built V-shaped holder and isolated from the immersion water of the objectives by an FEP film (50*µm*, *Adtech*). The translation of the sample through the light-sheet was performed by an actuator (*M-235, PI Instruments*). A *f* 7 = 200*mm* lens placed after the collection objective projects the image of the sample onto a camera (*Hamamatsu, Orca Flash 4.0*). The fluorescent response from the sample was filtered either by two filters with a sine/cosine transmission over the 400–700 nm range (*Optoprim*) as previously shown for a hyperspectral detection by Hedde et al.^26^ or by two bandpass filters for a conventional ratiometric detection (*Semrock FF02-447/60-25, FF01-560/94-25*) to collect NAD(P)H and FAD fluorescence, respectively. The filters were mounted on a fast filter wheel (*FW103H, Thorlabs*) to rapidly switch between them. Additional notch filters placed before the filter wheel were also used to block residual laser light scattered at 405 nm (405-13), 488 nm (488-15) if necessary. A permanent bandpass filter in the 400–700 nm range discards the light scattered at 375 nm whilst filtering the fluorescence. The synchronization between the lasers, the actuator, the filter wheel and the camera was achieved using MATLAB software.

### Data Acquisition

A total of *∼*120 embryos were imaged with *∼*10 embryos for each developmental stage per experimental replicate placed in the imaging holder. As only 2 embryos can be imaged simultaneously due to the dimension of the field of view, 5 volumetric acquisitions were performed per imaging session. The data for each imaging stack include approximately 30– 40 imaged planes, where each plane was imaged successively with different filters to recover different contrasts: i) no filter to collect the total intensity, ii) sine filter, iii) cosine filter, iv) Ch1 filter, v) Ch2 filter. The exposure time was 100 ms.

### Data processing

#### Data processing based on the phasor approach

The algorithm to process the metabolic data was developed in MATLAB. The intensity images acquired with no cosine/sine filter (*I_total_*), the sine (*I_sine_*) and cosine (*I_cosine_*) filters were first corrected from the background introduced by the ambient light in the lab room and the camera noise. The background references were chosen as the set of images from the first plane of the stack where the embryo is not visible in the field of the camera. Due to the use of different filters which may affect the background intensity detected by the camera, we decided to correct each image of the stack by the image from the first plane acquired with the same filter, i.e., image i of first plane to image i of plane N. *I_sine_* and *I_cosine_* images were then corrected by *I_total_* images so that they range from −1 to 1 using the following formula: 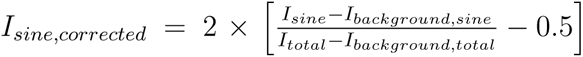, 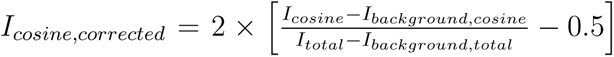. Data was then resized by a factor of 0.5 using bilinear interpolation. A region of interest around each individual embryo was manually selected from the image stack for further analysis. To consider only pixels containing relevant spectral information, we excluded pixels with intensities below a certain intensity threshold and then used the MATLAB *imerode* and *imdilate* operations to eliminate isolated pixels. The data were then plotted on a phase diagram using the *histogram2* function. The pair (*I_cosine,corrected_* and *I_sine,corrected_*) provided the abscissa and the ordinate coordinates, or equivalently (*I_modulus_*, *I_angle_*) for the modulus and the angular coordinates.

#### Definition of a RR for the hyperspectral approach

To assess changes in metabolic activity, we defined a metric by projecting the phasor coordinates of each image pixel along the axis joining the phasor coordinates of pure NAD(P)H and FAD species measured experimentally with the following formula:

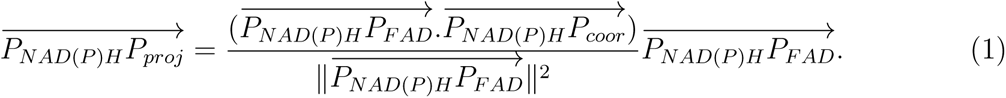

Then we calculated the relative contribution of the two metabolites in each pixel via the ratio: 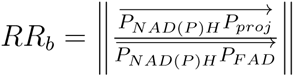. This value was encoded for each embryo.

The statistical study was performed by summing the histograms of RR data calculated for each imaged plans to extract a mean RR value for each embryo. The graphs presented in Figure 4.a describes the average values thus extracted for each embryo.

#### Definition of a RR for the conventional approach

To provide another metric against which to compare the previous metric, we estimated the RR via the conventional approach with bandpass filters. RR is derived from the following calculation: 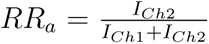 where *I_Ch_*_1_ is the intensity collected through the 447/60 filter and *I_Ch_*_2_ is the intensity collected through the 560/94 filter. Similarly to the hyperspectral approach, the average value of the RR histograms summed over all the imaged plane was extracted for every embryo.

### 3D reconstruction of hyperspectral data

For 3D reconstructions, a median filter was applied over three pixels to smooth the RR data. As the sample scan is performed along an axis tilted at 45°from the light-sheet axis, the recorded images and the RR data were transformed using affine transformation into physical coordinates. A linear interpolation (MATLAB *interp2*) was then performed on the intensity and RR volumes along the detection axis to achieve an isotropic pixel size. The two stacks were then centred and cropped to a cubic volume of the same size for all embryos. A mask was built from the intensity information and smoothed using a Gaussian function and then applied to the RR stack to remove low intensity pixels. Finally, RR values between −0.5 and 1 were encoded using a custom color map and IMARIS (Oxford Instruments, UK) software was used to obtain volumetric visualizations.

### Culture media preparation

All embryo culture took place in media overlaid with paraffin oil (*Merck Group, Darmstadt, Germany*) at 37°C in a humidified incubator set at 5% O_2_ and 6% CO_2_ balanced in N_2_. Culture dishes were pre-equilibrated for at least 4 h prior to use. All handling procedures were performed on microscopes fitted with heating stages calibrated to maintain media in dishes at 37°C. All culture media were supplemented with 4 mg/ml low fatty acid bovine serum albumin (*BSA, MP Biomedicals, AlbumiNZ, Auckland, NZ*) unless specified otherwise. Oviducts were collected in filtered Research Wash medium (*ART Lab Solutions, SA, Australia*) and embryos were cultured in filtered Research Cleave medium (*ART Lab Solutions, SA, Australia*).

### Embryos Preparation

Female (21-23 days) CBA x C57BL/6 first filial (CBAF1) generation mice were obtained from Laboratory Animal Services (*University of Adelaide, Australia*) and maintained on a 12 h light: 12 h dark cycle with rodent chow and water provided *ad libitum*. Animal ethics were approved by the School of Biology Ethics Committee of the University of St. Andrews (SEC20001) and the Animal Ethics Committee of the University of Adelaide (*M-2019-097*). Embryo collection was conducted in accordance with the Australian Code of Practice for the Care and Use of Animals for Scientific Purposes. Female mice were administered intraperitoneally (i.p.) with 5 IU of equine chorionic gonadotropin (*eCG; Folligon, Braeside, VIC, Australia*), followed by 5 IU human chorionic gonadotrophin (*hCG, i.p.; Kilsyth, VIC, Australia*) 46 h later. Female mice were then mated overnight with male mice of proven fertility. At 47 h post-hCG, females were culled by cervical dislocation and the oviducts were carefully dissected to isolate 2-cell embryos. Two-cell embryos were released from the oviducts by gently flushing the oviduct using pre-warmed Research Wash medium (*ART Lab Solutions, SA, Australia*) supplemented with 4 mg/ml low fatty acid bovine serum albumin (*BSA, MP Biomedicals, AlbumiNZ, Auckland, NZ*) using a 29-gauge insulin syringe with needle (*Terumo Australia Pty Ltd, Australia*) and subsequently underwent embryo vitrification.

### Embryo vitrification and warming

Media used for embryo vitrification and warming were as described in Tan et al.^14^ Briefly, two-cell embryos were vitrified with the Cryologic vitrification method, consisting of timely, sequential washes in handling medium followed by 3 mins in equilibration solution, and 30 s in vitrification solution, prior to loading onto a Fibreplug straw for storage in liquid nitrogen. For embryo warming, Fibreplugs containing embryos were removed from liquid nitrogen and quickly submerged into a handling medium supplemented with 0.3 M sucrose, followed by sequential washes in handling media with decreasing concentrations of sucrose (0.25, 0.15, and 0 M) for 5 min each. The post-warming survival rate was above 90% for all groups (data not shown). Warmed 2-cell embryos were then cultured in Research Cleave medium supplemented with 4mg/ml BSA and allowed to develop up until the blastocyst-stage.

### Sample preparation for imaging

For imaging, 2-, 4-, 8-cell, morula and blastocyst-stage embryos were collected at 6-, 10-, 24-, 30-, and 48-h post-warming respectively, and were transferred into pre-warmed 20 *µ*l drops of Research Wash medium supplemented with 4 mg/ml BSA in a specialised printed chamber overlaid with an FEP film and covered with paraffin oil. The Research Wash medium provides a physiologically pH-buffered medium for live embryo imaging. A maximum of 10 embryos per sample holder was used for imaging. Embryos were maintained at 37°C throughout the imaging process by heating the immersion water by means of two cartridges inserted in the support. The regulation of the voltage applied to the cartridges to keep the temperature constant is achieved with a PID controller.

## Acknowledgement

The authors would like to thank Mark Robertson for designing and building the objective support, Christopher Booth for designing and installing the sample mount heating system, and Juan Varela for providing the Imaris software.

## Supporting Information Available

The data that support the findings of the study are available from the corresponding author upon reasonable request.

