## Supplementary Material for "UVA hyperspectral light-sheet microscopy for volumetric metabolic imaging: application to pre-implantation embryo development"

### S1 - Determining the relationship between metabolic phasor coordinates, RR and the illumination wavelength

#### a) Theoretical relationship between RR and metabolic phasor coordinates

In this section, we investigate the relationship between metabolic phasor coordinates and the RR parameter. For this purpose, we consider the formal expression of the phasor coordinates (G, S) for a fluorescence signal arising from the co-excitation of the two co-enzymes, NAD(P)H and FAD. For example, the G phasor coordinate can be written as follows:

$$G = \frac{\int_{400}^{700} I^{\lambda_{exc}}(\lambda) T_{cos}(\lambda)\, d\lambda}{\int_{400}^{700} I^{\lambda_{exc}}(\lambda)\, d\lambda} \tag{1}$$

where $T_{cos}(\lambda)$ is the transmission of the cosine filter as a function of the wavelength. By substituting $I^{\lambda_{exc}}$ by $I^{\lambda_{exc}}(\lambda) = I^{\lambda_{exc}}_{NAD(P)H}(\lambda) + I^{\lambda_{exc}}_{FAD}(\lambda)$, the G phasor coordinate expression can be further expressed in terms of $G_{NAD(P)H}$ and $G_{FAD}$ as:

$$\begin{aligned} G = {} & \frac{\int_{400}^{700} I^{\lambda_{exc}}_{NAD(P)H}(\lambda)\, d\lambda}{\int_{400}^{700} \left( I^{\lambda_{exc}}_{NAD(P)H}(\lambda) + I^{\lambda_{exc}}_{FAD}(\lambda) \right) d\lambda} \frac{\int_{400}^{700} I^{\lambda_{exc}}_{NAD(P)H}(\lambda) T_{cos}(\lambda)\, d\lambda}{\int_{400}^{700} I^{\lambda_{exc}}_{NAD(P)H}(\lambda)\, d\lambda} \\ & + \frac{\int_{400}^{700} I^{\lambda_{exc}}_{FAD}(\lambda)\, d\lambda}{\int_{400}^{700} \left( I^{\lambda_{exc}}_{NAD(P)H}(\lambda) + I^{\lambda_{exc}}_{FAD}(\lambda) \right) d\lambda} \frac{\int_{400}^{700} I^{\lambda_{exc}}_{FAD}(\lambda) T_{cos}(\lambda)\, d\lambda}{\int_{400}^{700} I^{\lambda_{exc}}_{FAD}(\lambda)\, d\lambda} \\ & = \frac{I^{\lambda_{exc}}_{NAD(P)H}}{I^{\lambda_{exc}}_{NAD(P)H} + I^{\lambda_{exc}}_{FAD}} G_{NAD(P)H} + \frac{I^{\lambda_{exc}}_{FAD}}{I^{\lambda_{exc}}_{NAD(P)H} + I^{\lambda_{exc}}_{FAD}} G_{FAD} \end{aligned} \tag{2}$$

We can express this in terms of RR as:

$$G = (1 - RR) * G_{NAD(P)H} + RR * G_{FAD} \tag{3}$$

where $RR = \frac{I_{FAD}^{\lambda_{exc}}}{I_{NAD(P)H}^{\lambda_{exc}}+I_{FAD}^{\lambda_{exc}}}$ follows the original RR definition.[1] Similarly, an equivalent formula can be found for S with $S_{NAD(P)H}$ and $S_{FAD}$. Therefore, it is evident that the (G, S) phasor coordinates describe the position of a barycenter $P_{barycenter}$ between $P_{NAD(P)H}$ $(G_{NAD(P)H}, S_{NAD(P)H})$ and $P_{FAD}$ $(G_{FAD}, S_{FAD})$ where the weighting coefficients are $x_{NAD(P)H} = (1 - RR)$ and $x_{FAD} = RR$. This can be written as:

$$\overrightarrow{OP_{barycenter}} = (1 - RR)\,\overrightarrow{OP_{NAD(P)H}} + RR\,\overrightarrow{OP_{FAD}} \tag{4}$$

Specifically, the ratio between the distances $\left\|\overrightarrow{P_{NAD(P)H}P_{barycenter}}\right\|$ and $\left\|\overrightarrow{P_{NAD(P)H}P_{FAD}}\right\|$ is the RR value:

$$RR = \left\| \frac{\overrightarrow{P_{NAD(P)H}P_{barycenter}}}{\overrightarrow{P_{NAD(P)H}P_{FAD}}} \right\| \tag{5}$$

#### b) Theoretical relationship between RR, NAD(P)H-FAD concentration and illumination wavelength

Here, we develop further the relationship between RR and metabolite concentrations by deriving metabolic co-enzyme intensities using Beer Lambert's law. For example, $I_{NAD(P)H}^{\lambda_{exc}}$ can be written as:

$$I_{NAD(P)H}^{\lambda_{exc}}(\omega) \simeq log(10)\eta_{NAD(P)H}\alpha_{NAD(P)H}^{\lambda_{exc}}N_{NAD(P)H}M\hbar\omega p_{NAD(P)H}(\omega) \tag{6}$$

where $\eta_{NAD(P)H}$ is the quantum yield of NAD(P)H molecule, $\alpha_{NAD(P)H}^{\lambda_{exc}}$ its molar absorbance at $\lambda_{exc}$ excitation wavelength, $N_{NAD(P)H}$ the number of molars per unit volume, $M$ the number of incident photons and $p_{NAD(P)H}$ is the probability density function in the spectral domain.

When collecting the intensity over the spectral range $[\omega_1; \omega_2]$, the intensity can be rewritten as:

$$I_{NAD(P)H}^{\lambda_{exc}} \simeq log(10)\eta_{NAD(P)H}\alpha_{NAD(P)H}^{\lambda_{exc}} N_{NAD(P)H} M\hbar\omega IP_{NAD} \quad (7)$$

where $IP_{NAD} = \int_{\omega_1}^{\omega_2} p_{NAD(P)H}(\omega)\, d\omega$. As result, the weighting coefficients $\{x_{NAD(P)H}; x_{FAD}\}$ can be rewritten as:

$$\begin{cases} x_{NAD(P)H} = \frac{\eta_{NAD(P)H}\alpha_{NAD(P)H}^{\lambda_{exc}} N_{NAD(P)H} IP_{NAD(P)H}}{\left(\eta_{NAD(P)H}\alpha_{NAD(P)H}^{\lambda_{exc}} N_{NAD(P)H} IP_{NAD(P)H} + \eta_{FAD}\alpha_{FAD}^{\lambda_{exc}} N_{FAD} IP_{FAD}\right)} \\ x_{FAD} = \frac{\eta_{FAD}\alpha_{FAD}^{\lambda_{exc}} N_{FAD} IP_{FAD}}{\left(\eta_{NAD(P)H}\alpha_{NAD(P)H}^{\lambda_{exc}} N_{NAD(P)H} IP_{NAD(P)H} + \eta_{FAD}\alpha_{FAD}^{\lambda_{exc}} N_{FAD} IP_{FAD}\right)} \end{cases} \quad (8)$$

or in a simple form as:

$$\begin{cases} x_{NAD(P)H} = \frac{\frac{a}{b} N_{NAD(P)H}}{\frac{a}{b} N_{NAD(P)H} + N_{FAD}} \\ x_{FAD} = \frac{N_{FAD}}{\frac{a}{b} N_{NAD(P)H} + N_{FAD}} \end{cases} \quad (9)$$

where $a = \eta_{NAD(P)H}\alpha_{NAD(P)H}^{\lambda_{exc}} IP_{NAD(P)H}$ and $b = \eta_{FAD}\alpha_{FAD}^{\lambda_{exc}} IP_{FAD}$. Therefore, the weighting coefficient $x_{FAD}$ correctly reflects the concentration ratio $N_{FAD}/(N_{NAD(P)H} + N_{FAD})$ provided that $a = b$, i.e. more specifically, if the following equation is satisfied:

$$\eta_{NAD(P)H}\alpha_{NAD(P)H}^{\lambda_{exc}} IP_{NAD(P)H} = \eta_{FAD}\alpha_{FAD}^{\lambda_{exc}} IP_{FAD} \quad (10)$$

In the literature, quantum yield values for the two species have been reported to be equal to 0.019 (ref.[2]) and 0.033 (ref.[3]) for NAD(P)H and FAD respectively. The integral of the probability density function $\{IP_{NAD(P)H}, IP_{FAD}\}$ can be approximated to 1 when the detection spectral range is wide enough to completely capture NAD(P)H and FAD fluorescent emissions. Unlike quantum yield and probability density functions, molar absorbance values depend on the excitation wavelength. Based on the absorbance graphs available in the literature, it can be seen that only an excitation in the UVA range can achieve relatively similar absorbance values for NAD(P)H and FAD. As a conclusion, RR values derived from the weighting coefficient $x_{FAD}$ approximates the concentration ratio $N_{FAD}/(N_{NAD(P)H} + N_{FAD})$ for illumination in the UVA range, such that FAD and NAD(P)H are co-excited, simultaneously collected and in the absence of additional normalization.

#### c) Numerical comparison between RR assessment with bandpass filters and cosine/sine filters

Finally, we compared the accuracy and precision of RR values using the bandpass filtering approach and the spectral phasor analysis. Figure S1a shows normalized emission curves for pure NAD(P)H and FAD solutions, together with the emission curves for diluted solutions with a 10% dilution factor step between the pure solutions. Noise was added to these curves to simulate actual conditions ($SNR = 20$) and quantify values accuracy, as shown in Figure S1b.

Figure S1e shows RR values assessed with conventional filtering using the transmission curves indicated in Figure S1c, where narrow-band transmissions are displayed in black and large-band transmissions in grey. RR values - assessed via the ratio $RR = I_{Ch2}/(I_{Ch1} + I_{Ch2})$ - range from 0.5 and 1 when the dilution coefficient varies between 0 (pure NAD(P)H) and 1 (pure FAD). Average RR values are found to be similar for wide-band and narrow-band filters. To estimate the precision of this analysis, we calculated the ratio of the standard deviation of RR values divided by the associated gradient. The graph of this metric is shown in the Figure S1f, and indicates greater accuracy for wide-band filters compared to narrow-band filters due to the larger number of photons collected.

We then compared the accuracy and precision of RR derived from bandpass filtering to RR from the spectral phasor analysis. Prior to RR calculation, the phasor coordinates of the studied solutions are calculated and plotted in Figure S1d. Due to the normalization of the spectra over the 400-700nm range, the phasor coordinates are aligned and evenly distributed along the NAD(P)H-FAD trajectory. Figure S1e shows RR values derived from the formula

$RR = \left\| \frac{\overrightarrow{P_{NAD(P)H}P_{barycenter}}}{\overrightarrow{P_{NAD(P)H}P_{FAD}}} \right\|$. RR values follow a linear variation between 0 and 1 when the dilution coefficient varies between 0 and 1. These values show a high degree of accuracy compared to the true relation between RR values and the dilution factor represented with a dotted linear line. Figure S1f also shows how the precision of the spectral phasor analysis compares with the conventional filtering. Interestingly, we found better precision for the phasor analysis compared to filtering with narrow-band transmission, and to a lower extent for the filters with wide-band transmission.

To better understand the effect of the normalization of the emission spectra on the linearity of the RR parameter, we considered emission profiles where the normalization factor of the pure FAD solution is 10 times higher than that of the NAD(P)H solution. Figure S1g shows that the phasor coordinates are no more evenly distributed along the FAD-NAD(P)H trajectory, although the average phasor coordinate of pure NAD(P)H and FAD is only slightly affected. As shown in Figure S1h, this means that the spectral phasor analysis of RR values does not vary linearly with the dilution coefficient, as is the case for the RR evaluated with the two bandpass filters. Interestingly, RR values derived from phasor analysis maintain a slightly more accurate trend compared to RR values derived from bandpass transmission. Figure S1i shows lower precision at low dilution coefficients for both types of filtering. Phasor analysis provides slightly higher precision than bandpass filtering for wide spectral bandwidths, whereas a great loss of precision occurs with small spectral bandwidths. This last case, where the spectra are not normalized, can be considered as a case where the equality of equation 10 is not verified, with, for example, a very different absorbance between NAD(P)H and FAD, showing the importance of meeting the equality to maintain the linearity of the RR with concentration.

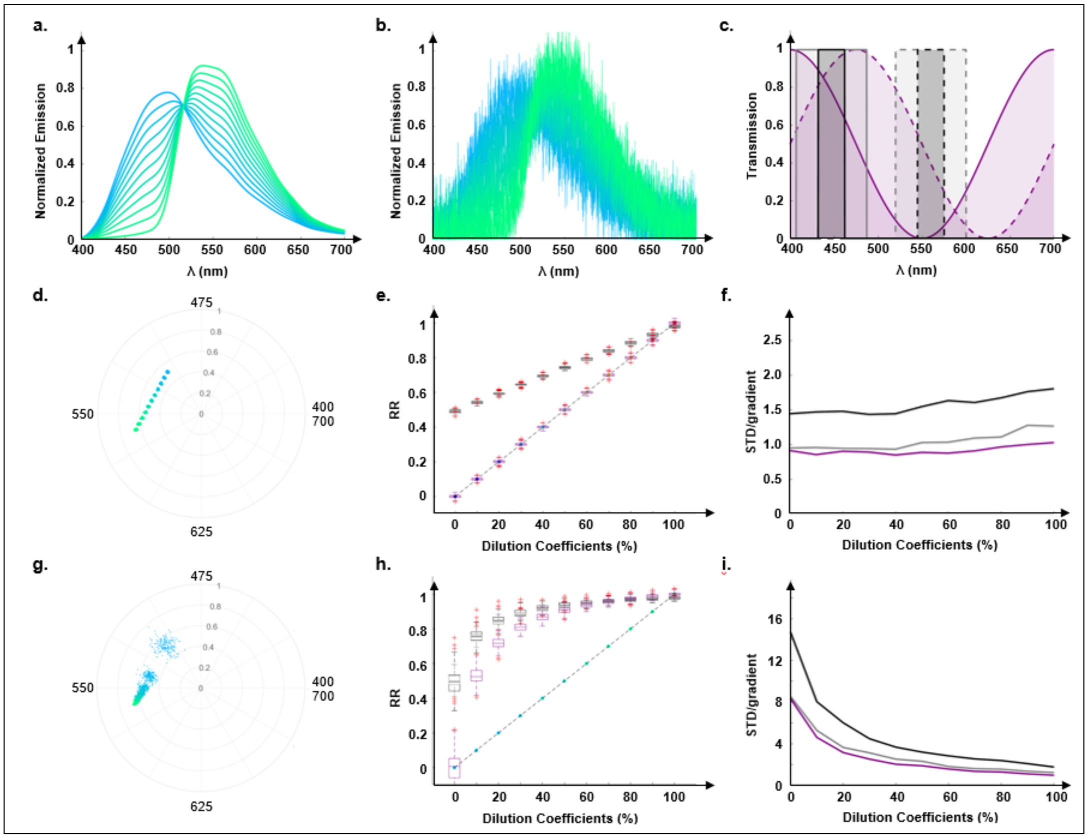


Figure S1: (a) Emission spectra of the dilutions between NAD(P)H and FAD solutions (blue: pure NAD(P)H, green: pure FAD). (b) Emission spectra with additional noise, SNR = 20. (c) Transmission profiles of cosine-sine (purple) and bandpass filters (black: thin bandwidth, grey: large bandwidth). (d) Spectral phasor plot. (e) Graphs of RR values calculated using spectral phasor method (purple) and bandpass filtering (grey, black). (f) Graphs of RR standard deviation normalized by RR gradient for both approaches. (g) Spectral phasor plot where NAD(P)H pure solution intensity exceeds FAD pure solution intensity by a factor of 10. (h) Graph of RR values and (i) Graph of RR standard deviation normalized by RR gradient in this case.

### S2 - Setup

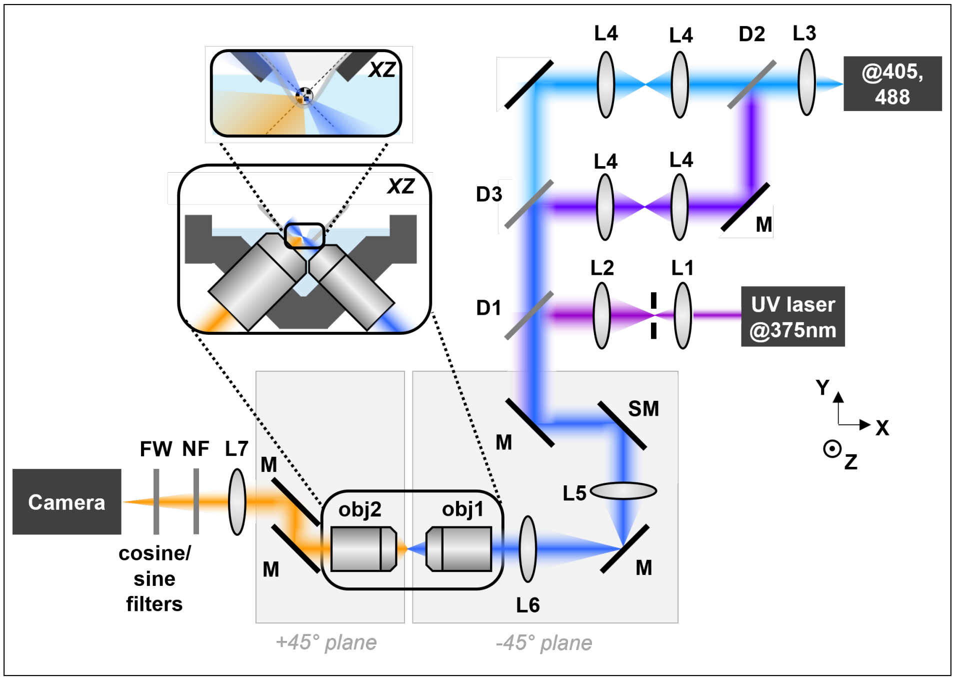


Figure S2: Experimental setup. The 375-nm UV beam is expanded by a telescope formed by $\{L1; L2\}$ then uplifted by a periscope (P); the other two beams at 405 and 488 nm are split using a dichroic mirror (D2) and relayed individually through 1:1 telescopes (L4) before being uplifted by two periscopes (P) I dont see a periscope in the fig. I dont think the exact implementation needs to be mentioned.. The 405 and 488-nm beams are then recombined with a dichroic mirror (D3), and recombined with the 375-nm UV beam with another dichroic mirror (D1). The overlapped beams are sent onto a scanning mirror (SM) conjugated to the back pupil plane of the illumination objective (obj1) by means of a telescope $\{L5; L6\}$. Fluorescent light is collected by the detection objective (obj2) and detected onto the camera via a lens (L7). Filters are mounted in a filter wheel (FW) and notch filters (NF) are placed in the detection path to remove scattered light. The sample is mounted in a V-shaped sample holder, illustrated in the inset.

### S3 - Results on the safety of lightsheet imaging at 375 nm

Embryos at each developmental stage were subjected to imaging with our 375 nm hyperspectral light sheet and were then returned to the incubator immediately for subsequent culture up to the blastocyst stage. As shown in Figure S3, we found that more than 70% of embryos developed to the blastocyst stage following imaging, which is similar to the developmental rate of non-imaged (control) embryos, indicating that exposure to our 375 nm hyperspectral light sheet does not negatively impact embryo development.

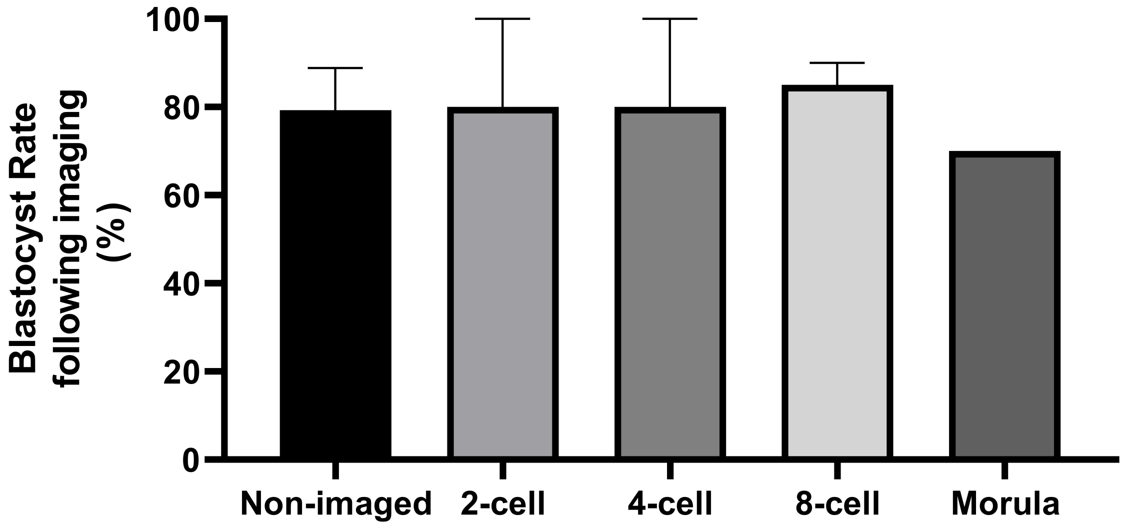


Figure S3: Exposure to 375nm during imaging is safe for embryo development. Embryos were exposed to 375nm laser during imaging on the hyperspectral light sheet system and were returned to the incubator for subsequent culture up to the blastocyst stage. Data are presented as mean ± SEM, n = 20-32 embryos per developmental stage, from 2 independent experimental replicates.

#### S4 - Correlation between hyperspectral and conventional methods

To determine whether changes in embryo RR using the hyperspectral method truly correlate to the RR estimated with the conventional filters method, we performed correlative analysis of the RR obtained from the two methods. The scatter plots of RR of filters versus hyperspectral method for each embryo across all developmental stages depict a strong positive correlation between the two methods (cf. Figure S4; $P<0.05$). Collectively, these results confirm the validity of our hyperspectral method for detecting changes in RR during preimplantation embryo development.

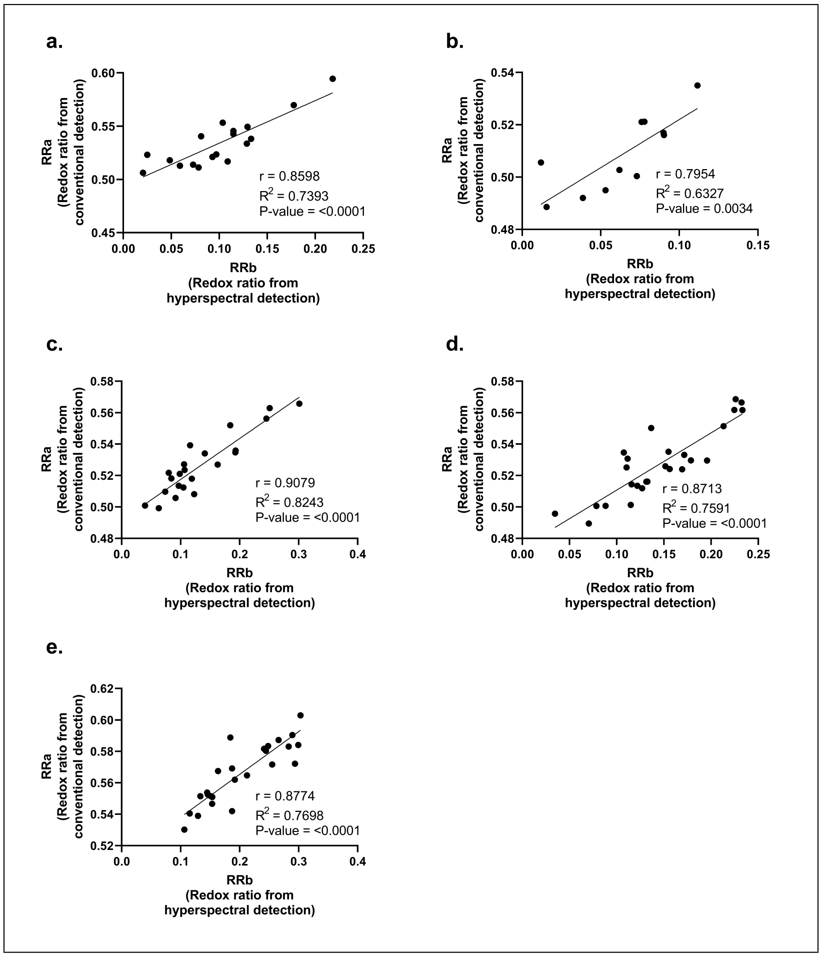


Figure S4: Scatter plots of the computed redox ratio with conventional and phase-based hyperspectral collection across different developmental stages. Embryos throughout development were subjected to imaging with our 375 nm hyperspectral light sheet. Specifically, the embryo developmental stages imaged were (a) 2-cell; (b) 4-cells; (c) 8-cells; (d) morula- and (e) blastocyst-stage. The fluorescence signals were collected either via the conventional method with two specific bandpass filters or the phase-based hyperspectral approach. Results of linear regression analysis (represented as a black solid line) and the computed Pearson correlation ($r$) values indicate a strong positive relationship between the computed redox ratio using the two methods. P-values<0.001 for all stages of development. n = 11-26 embryos for each developmental stage, from 3 independent experimental replicates.

(2) Scott, T. G.; Spencer, R. D.; Leonard, N. J.; Weber, G. Emission properties of NADH.

Studies of fluorescence lifetimes and quantum efficiencies of NADH, AcPyADH, and simplified synthetic models. *J. Am. Chem. Soc* **1970**, *92*, 687–695.

(3) Islam, S. D.; Susdorf, T.; Penzkofer, A.; Hegemann, P. Fluorescence quenching of flavin adenine dinucleotide in aqueous solution by pH dependent isomerisation and photo-induced electron transfer. *Chemical physics* **2003**, *295*, 137–149.
